# Sub-Additive Effects of Autism-ADHD Comorbidity on Resting-State Pupillary and Oculomotor Phenotypes

**DOI:** 10.64898/2026.05.27.728181

**Authors:** Xin Di, Bharat B. Biswal

## Abstract

**Background:** Autism Spectrum Disorder (ASD) and Attention-Deficit/Hyperactivity Disorder (ADHD) share substantial clinical and physiological overlap. While naturalistic and sensory-driven paradigms increasingly capture evoked neurophysiological responses, the intrinsic baseline physiology of these conditions remains poorly defined. We characterized resting-state pupillary volatility and oculomotor stability across the ASD-ADHD spectrum using dimensional and categorical (DSM-5) frameworks.

**Methods:** We analyzed resting-state eye-tracking data from a large pediatric cohort (N = 2,315) from the Healthy Brain Network, extracting Pupil Relative Volatility (Coefficient of Variation [CV]) and Bivariate Contour Ellipse Area (BCEA) to index pupillary and spatial gaze stability. Data were evaluated via continuous regressions against Social Responsiveness Scale (SRS) and SWAN inventories, then 2×2 factorial ANCOVAs based on clinical diagnoses, with sensitivity analyses for extreme values, hardware heterogeneity, and psychostimulant medication.

**Results:** Dimensional models revealed no significant association between either metric and continuous ASD or ADHD trait severity, apart from a modest sub-additive SRS × SWAN interaction on pupillary CV; an apparent pooled BCEA-trait association proved attributable to hardware differences and was absent within hardware-consistent subgroups. Categorical models showed robust diagnostic differentiation for both metrics: isolated ASD was associated with elevated CV and impaired BCEA, with ADHD additionally impairing BCEA. Comorbid ASD+ADHD produced a sub-additive CV interaction robust across outlier-resistant, hardware-stratified, and medication-adjusted analyses; an analogous BCEA interaction was directionally consistent but not significant under standard estimation.

**Conclusions:** Categorical diagnostic status was consistently associated with baseline pupillary and oculomotor physiology across the ASD-ADHD spectrum in this cohort, with a shared sub-additive comorbidity effect most robustly expressed in pupillary volatility; continuous trait severity showed little independent association with either metric. These findings, obtained using a rigorously validated, hardware-heterogeneous multi-site sample, establish a foundation for future naturalistic and sensory-evoked investigations of the shared physiological architecture underlying ASD-ADHD comorbidity.

## 1. Introduction

Autism Spectrum Disorder (ASD) and Attention-Deficit/Hyperactivity Disorder (ADHD) are highly prevalent neurodevelopmental conditions characterized by substantial clinical and biological overlap; modern epidemiological estimates indicate that up to 70% of autistic youths also meet criteria for ADHD (Lai et al., 2019; Ronald et al., 2008). Despite shared phenotypic presentations, including executive dysfunction, motor atypicalities, and sensory dysregulation, traditional diagnostic frameworks such as the DSM-5 rely on categorical thresholds that may mask continuous biological variance underlying these conditions. The Research Domain Criteria (RDoC) initiative (Cuthbert and Insel, 2013) has motivated efforts to map continuous symptom dimensions, autism traits via the Social Responsiveness Scale (SRS; (Constantino, 2021)) and attention-deficit traits via the SWAN scale (Swanson et al., 2012), onto physiological measures in search of transdiagnostic mechanisms, though whether categorical or dimensional frameworks better characterize any given physiological measure is an empirical question rather than a foregone conclusion.

Resting-state paradigms, which record activity in the absence of explicit cognitive or stimulus-driven demands, offer a window into endogenous autonomic tone and oculomotor stability distinct from what task-based and naturalistic viewing paradigms capture (Aminihajibashi et al., 2019; Di Martino et al., 2014; Kim and Alvarez, 2013). Spontaneous pupillary fluctuation during rest has been proposed as a peripheral correlate of autonomic arousal, with theoretical links to Locus Coeruleus-Norepinephrine (LC-NE) activity classically established for the phasic, stimulus-evoked pupillary response (Aston-Jones and Cohen, 2005; Joshi et al., 2016); the neurophysiological substrates of tonic, resting-state pupillary variability specifically are less well characterized. Existing evidence on resting-state autonomic function in autism is mixed: a systematic review of 60 studies found evidence of group differences in only 61% of comparisons, with inconsistent direction across studies and outcome measures (Arora et al., 2021), and direct pupillometric findings range from larger baseline pupil size in ASD (Anderson et al., 2013; Anderson and Colombo, 2009) to null or reversed effects (Nuske et al., 2016). Trait-linked pupillary adaptation has also been reported to scale continuously with autism symptom severity independent of categorical diagnosis (DiCriscio and Troiani, 2017), consistent with a dimensional account, though this literature has not, to our knowledge, been extended to ADHD or to comorbid presentations. Task-evoked studies further support a role for dysregulated autonomic arousal in shaping oculomotor performance in ASD and ADHD, with pre-saccadic pupil size differentially associated with saccadic reaction time by diagnosis, linearly in autism, quadratically in ADHD (Bellato et al., 2023), though whether comparable arousal-oculomotor coupling exists at baseline, independent of any task, sensory stimulus, or attentional demand, has not been established. Non-additive effects of ASD and ADHD on other oculomotor and social-attentional measures have also been reported in a comorbid sample (Seernani et al., 2021), consistent with our hypothesis that comorbidity produces a distinct rather than purely cumulative physiological profile.

Gaze stability during fixation similarly indexes cortico-cerebellar and brainstem oculomotor control, and has been linked to attentional and motor dysregulation in neurodevelopmental populations (Munoz et al., 2003). The Bivariate Contour Ellipse Area (BCEA) provides a two-dimensional index of fixational gaze dispersion that improves on independent 1D coordinate analyses (Crossland and Rubin, 2002; Timberlake et al., 2005) and has recently been applied to ADHD directly, with children with ADHD showing significantly greater BCEA than typically developing peers during a sustained-fixation task (Moiroud et al., 2025); to our knowledge, this measure has not previously been examined dimensionally against continuous ADHD or autism trait severity, nor in the context of ASD-ADHD comorbidity.

To investigate the shared and distinct intrinsic physiological baselines of the ASD-ADHD spectrum, we analyzed unconstrained resting-state eye-tracking data from a large pediatric cohort within the Healthy Brain Network (HBN). We extracted Pupil Relative Volatility (CV) and Spatial Gaze Instability (BCEA) and evaluated both metrics using dimensional (continuous trait) and categorical (DSM-5 diagnosis) analytical frameworks in parallel, rather than assuming a priori which framework would best characterize either measure. We hypothesized that comorbid ASD+ADHD would be associated with a distinct, non-additive physiological signature relative to either condition in isolation, a sub-additive rather than simply cumulative pattern, reflecting an interactive rather than purely additive relationship between the two conditions’ effects on baseline oculomotor and pupillary physiology.

## 2. Materials and Methods

### 2.1. Study Population and Clinical Characterization

Data were obtained from the HBN sample, a large-scale open-science initiative generating a transdiagnostic pediatric psychiatric dataset via community-referred recruitment based on perceived clinical concern rather than epidemiologic ascertainment (Alexander et al., 2017). Inclusion required an estimated IQ ≥ 70 and the absence of acute safety concerns or medical conditions expected to confound brain-related findings. The initial dataset comprised all participants who completed the resting-state eye-tracking paradigm; following quality control (block-level gaze-completeness thresholds), the final analytic sample consisted of 2,315 children and adolescents. Pupil-channel calibration validity was evaluated independently of this inclusion criterion; 1,448 participants (62.5%) additionally had a usable calibrated pupil signal and were eligible for pupillary CV-based analyses, with sample sizes reported separately for pupil- and gaze-based models throughout. Missingness reflected technical attrition and task non-compliance during recording rather than clinical dropout or missing phenotypic data.

Categorical diagnoses were determined by research clinicians using the computerized Kiddie Schedule for Affective Disorders and Schizophrenia (KSADS-COMP), cross-referenced with parent-report instruments and behavioral observations to establish DSM-5 consensus. Categorical models stratified participants into four groups: Typically Developing controls (TD; no consensus diagnosis), isolated ASD, isolated ADHD, and Comorbid (ASD+ADHD). Participants with other primary psychiatric conditions (e.g., primary anxiety or mood disorders) without concurrent ASD/ADHD were retained for dimensional models but excluded from categorical analyses to prevent phenotypic confounding. Stimulant medication use at the time of testing was recorded via parent report for sensitivity analyses (Supplementary Section S3).

Continuous trait severity was assessed via two parent-report inventories, each subject to its own availability. Autistic trait severity was quantified via the SRS-2 Total T-score, standardized to allow merging of the preschool and school-age forms administered across this age range. Inattentive/hyperactive-impulsive trait severity was measured via the SWAN Total score. Both inventories were Z-scored within each model’s analytic sample prior to modeling.

### 2.2. Resting-State Paradigm

Participants underwent a resting-state paradigm in a sound-shielded room, seated facing a computer display at a fixed viewing distance; exact screen geometry and viewing distance varied by acquisition site and are addressed directly in Sections 2.3–2.4. The paradigm was designed to capture intrinsic, endogenous physiological and cortical baseline activity in the absence of external cognitive or emotional stimuli.

A central fixation cross was displayed throughout, with participants auditorily instructed to open or close their eyes at alternating intervals: five 20-second “eyes-open” blocks alternating with five 40-second “eyes-closed” blocks. Only data from the eyes-open blocks were used in the present oculomotor and pupillary analyses.

### 2.3. Eye-Tracking and Pupillometry Acquisition

Simultaneous EEG was recorded but is not analyzed here. Published HBN protocol documentation describes a single infrared eye tracker (iView-X RED-m, SensoMotoric Instruments GmbH) operating at 120 Hz, with 0.1° spatial resolution and 0.5° gaze accuracy. Inspection of the raw files underlying this sample, however, showed recordings distributed across three nominal sampling rates (30, 60, and 120 Hz) that differed systematically in acquisition characteristics beyond rate: the 30/60 Hz configurations included corneal-reflection (CR)-based pupil correction and a calibrated diameter channel (mm), while the 120 Hz configuration did not, providing only an uncorrected pixel-space measurement. We could not fully reconcile this with the single-system protocol description using public sources; it may reflect an equipment, firmware, or export-software change over this ongoing, multi-site collection effort postdating the original protocol publication. We treated hardware configuration and sampling rate as empirical properties of each recording to be directly assessed and corrected for (Section 2.4), rather than assuming acquisition uniformity.

A standardized 5-point calibration protocol was administered prior to data collection, repeated until spatial error was <2° for any single point and <1° on average, as documented for the SMI system.

### 2.4. Eye-Tracking and Pupillometry Preprocessing

Raw data were preprocessed via a custom Python pipeline isolating valid physiological signals during “eyes-open” resting-state blocks. Gaze and pupil time series were segmented into 20-second epochs based on hardware triggers. The first 3 seconds of each epoch were excluded to avoid contamination by the pupillary light reflex, a luminance-driven transient distinct from the intrinsic fluctuations of interest (Di et al., 2026; Mathôt et al., 2018), yielding a ∼17-second analysis window per block.

Each raw file was inspected for a calibrated pupil-diameter channel (mm, corneal-reflection [CR]-derived). One of three tracker configurations lacked CR correction and provided only an uncorrected pixel-space measurement, confounded by head-to-camera distance in a manner that varies within-recording and cannot be resolved via linear rescaling; files without a calibrated channel were excluded from pupil-based analyses (62.5% of the eligible sample retained a valid channel), determined per-file rather than by sampling rate. Gaze-based analyses do not depend on pupil validity and retained the full eligible sample.

Blinks and tracking loss were flagged via zero/missing values. Where a calibrated pupil channel existed, samples were additionally screened for physiologically implausible dilation speed and trend-line deviation (each: median + 4.5 MADs (Kret and Sjak-Shie, 2019)), with a 75 ms rejection margin applied around each flagged sample to account for peri-artifact contamination. This pupil-artifact mask was not propagated to gaze, so pupil-specific artifacts do not discard valid gaze data; genuine blinks (zero-value pupil, pre-flagging) were treated as invalidating simultaneous gaze position, since gaze cannot be measured with the eye closed.

Gaps ≤500 ms were linearly interpolated per channel. Blocks with <19 s of active tracking (pre-trim) were discarded outright; among retained blocks, >20% post-interpolation missingness in the gaze channels (not pupil) determined exclusion, so blocks with valid gaze but unusable pupil data remain gaze-eligible. Participants required ≥2 valid blocks (range 2–5), included as a covariate in all models. For pupil-valid participants, the per-block flagged-sample proportion (blink, artifact, margin) was averaged across blocks as a data-quality covariate (percent artifact) in pupil-CV models only, since it is undefined without a pupil channel and not applicable to gaze outcomes.

Raw gaze position (pixels) was converted to degrees of visual angle using each file’s own calibration geometry (screen resolution, physical stimulus dimensions) and viewing distance, since these varied by recording session rather than uniformly by tracker, placing gaze on a common, hardware-independent scale. All time series (30/60/120 Hz native rates) were downsampled to 30 Hz via mean-bin resampling.

### 2.5. Feature Extraction: Pupillary Volatility and Spatial Gaze Instability

Unlike naturalistic viewing paradigms that assess stimulus-driven entrainment, resting-state paradigms measure intrinsic, endogenous physiological fluctuations. We extracted two metrics to quantify baseline pupillary volatility and oculomotor stability, each computed independently for every valid block and then averaged across a participant’s valid blocks to yield a single, stable trait-level estimate. This block-level approach avoids conflating genuine within-block fluctuation with between-block shifts in signal level (e.g., due to habituation, recording drift, or head repositioning), which would otherwise inflate a pooled, across-block estimate independent of any true physiological or oculomotor variability.

*Pupillary Relative Volatility (Coefficient of Variation).* Absolute baseline pupil size is confounded by individual physiological differences and age-related miosis. To obtain a standardized, unit-invariant index of endogenous pupillary fluctuation, we computed the coefficient of variation (CV = σ/μ) for pupil diameter, using the artifact-cleaned, PLR-trimmed calibrated pupillary time series. Tonic pupillary CV of this kind has been proposed as an index of autonomic/arousal-related fluctuation; however, the neurophysiological substrates of resting-state pupillary variability are less well characterized than those of the phasic, stimulus-evoked pupillary response classically linked to the LC-NE system, and pupillary CV is treated here as a candidate physiological correlate rather than a validated marker of a specific neural mechanism.

*Spatial Gaze Instability (BCEA).* Gaze position stability during the eyes-open baseline was quantified using the BCEA, computed from the horizontal and vertical gaze coordinates (in degrees of visual angle) as

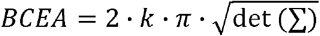

where Σ is the 2×2 covariance matrix of horizontal and vertical gaze position and k = −ln(1 − P), with P = 0.683 corresponding to a standard 1-σ bounding ellipse. BCEA is reported in square degrees of visual angle (deg²).

Because pupil-channel validity and gaze-channel validity are independent properties of a given recording, the analytic samples for pupil-CV-based and BCEA-based models differ: BCEA analyses draw on the full gaze-eligible sample, while pupil-CV analyses are additionally restricted to participants with a valid calibrated pupil channel. Sample sizes are reported separately for each analysis throughout.

### 2.6. Statistical Analysis

All modeling was performed in Python (statsmodels). All models controlled for participant age, biological sex, hardware sampling rate, and valid resting-state block count (2–5) as baseline covariates. Because pupil- and gaze-channel validity are independent properties of each recording, pupillary CV and BCEA models were fit on separate analytic samples via metric-specific listwise deletion rather than a shared requirement. CV models additionally included the pre-interpolation flagged-sample rate (percent artifact) as a covariate, omitted from BCEA models as this metric is undefined without a pupil signal. Sample sizes are reported separately by metric throughout. Benjamini-Hochberg false-discovery rate (FDR) correction was applied within each physiological metric family.

#### 2.6.1. Dimensional Modeling of Continuous Traits

Physiological outcomes (Pupil CV, BCEA) and continuous predictors (SRS-2, SWAN) were Z-scored within each metric’s analytic sample for standardized OLS coefficients (β), fit in three stages: (1) *Separate* models evaluating SRS and SWAN independently; (2) *Joint* models evaluating both simultaneously to isolate trait-specific variance from their behavioral covariance; and (3) *Interaction* models (SRS × SWAN) testing whether co-occurring elevated traits relate non-linearly to baseline physiology. Effect sizes were quantified as incremental R² (ΔR²) over the baseline covariate model.

#### 2.6.2. Categorical (DSM-5) Threshold Validation

Participants with heterogeneous (“Other”) clinical presentations were excluded, and the remaining cohort stratified into a 2×2 factorial design (ASD Status × ADHD Status), with typically developing controls as baseline reference. A 2×2 ANCOVA evaluated main diagnostic effects and their interaction (partial η²), testing whether comorbid ASD+ADHD relates non-additively to baseline physiology relative to either diagnosis alone.

#### 2.6.3. Sensitivity Analyses

Three sensitivity analyses assessed stability against methodological, hardware, and pharmacological confounds (Supplementary Materials): 1) *Outlier Resistance* (Section S1) re-fit the categorical models via robust linear modeling (Huber’s T) to test dependence on high-leverage observations. 2) *Hardware Subgroup Sensitivity* (Section S2) re-evaluated both frameworks restricted to hardware-consistent recordings, given documented calibration and gaze-processing differences across tracker configurations. The only check to alter a substantive conclusion, as the pooled dimensional BCEA-trait association did not survive restriction and is not interpreted as genuine, whereas categorical main effects and the pupillary CV interaction were unaffected. And 3) *Psychostimulant Medication* (Section S3) characterized medication-log missingness and prevalence, then re-evaluated categorical models with active stimulant use as an explicit covariate.

## 3. Results

### 3.1. Participant Demographics and Data Quality

The final analyzed resting-state cohort comprised 2,315 pediatric participants (1,452 males, 863 females; mean age 11.05 years, SD = 3.49), with a broad distribution of trait severities across the full transdiagnostic sample (mean SRS-2 T-score 57.33, SD = 11.51; mean SWAN total 0.46, SD = 1.00; Table 1).

**Table 1.**
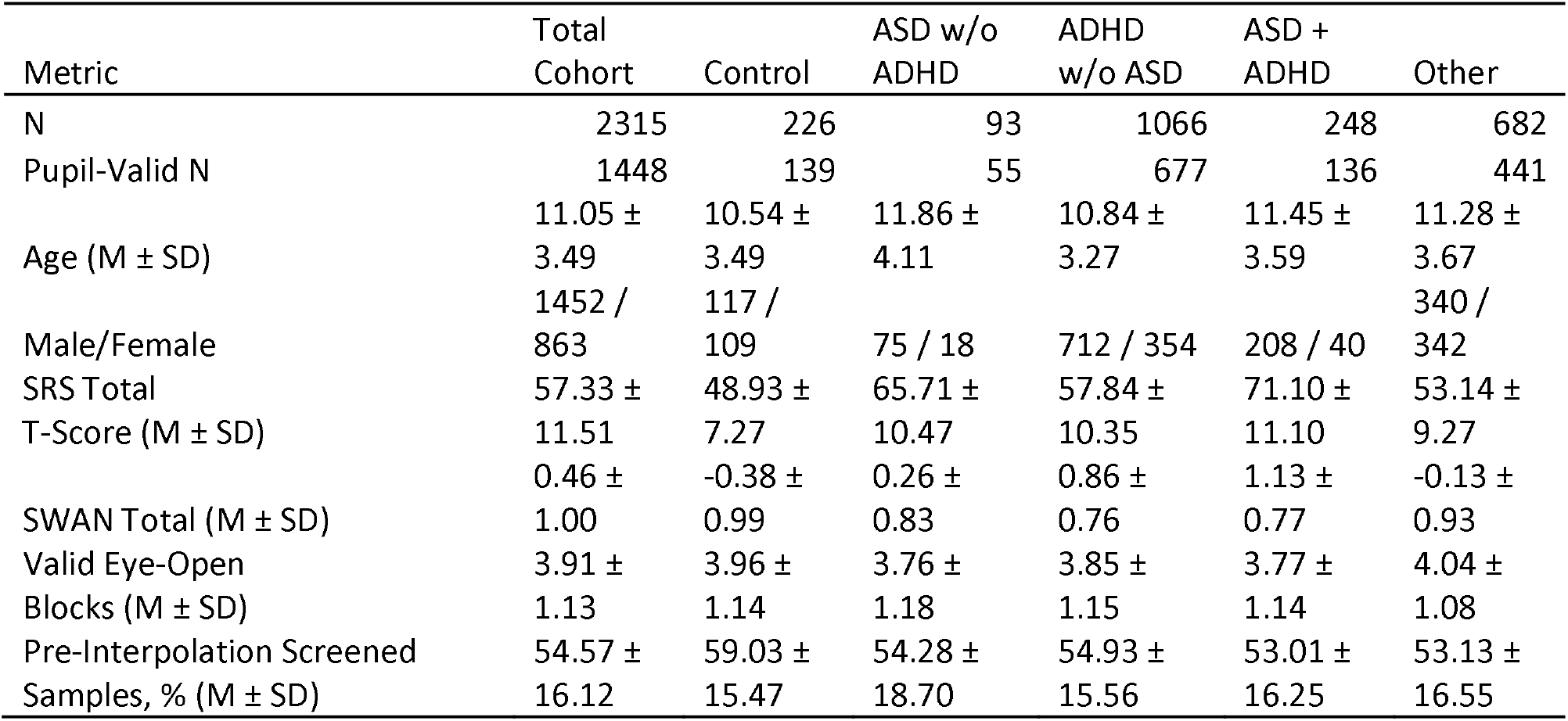
Demographic and clinical characteristics of the resting-state cohort (mean ± SD). ‘Other’ includes participants with psychiatric/neurodevelopmental diagnoses outside the primary ASD/ADHD groups, retained only in dimensional models. SRS-2 and SWAN scores were missing for 43 and 57 participants, respectively; reported means reflect available data. All participants are eligible for BCEA-based analyses; Pupil-Valid N indicates the subset eligible for CV-based analyses (62.5% of the cohort; Section 2.4). Pre-Interpolation Screened Samples, % is computed among pupil-valid participants only and reflects blink, statistical-artifact, and edge-margin flagging prior to interpolation; most flagged intervals were recovered by interpolation and this figure does not represent unrecoverable data loss (Section 2.4).

Clinical subgroups comprised 1,066 children with isolated ADHD, 93 with isolated ASD, 248 with comorbid ASD+ADHD, and 226 typically developing controls; an additional 682 participants with other psychiatric or neurodevelopmental diagnoses were retained for dimensional but not categorical models.

All 2,315 participants were eligible for BCEA-based analyses; 1,448 (62.5%) additionally had a usable pupil signal and were eligible for CV-based analyses, with sample sizes reported separately by metric throughout.

Participants contributed an average of 3.91 valid blocks (SD = 1.13), included as a covariate in all models. Among pupil-valid participants, an average of 54.6% (SD = 16.1) of samples per block were flagged prior to interpolation (percent artifact; Table 1, Section 2.4), included as a covariate in pupil-CV models only. Full demographic characteristics by diagnostic group are in Table 1.

### 3.2. Dimensional (RDoC) Analysis of Continuous Traits

Continuous trait associations were assessed via separate, joint, and interaction models for SRS-2 and SWAN scores (Table 2), with pupillary CV evaluated in the pupil-valid subset (N = 1,408) and BCEA across the full gaze-eligible cohort (N = 2,238).

**Table 2.** Dimensional regression models of resting-state physiology on ASD/ADHD trait severity. Pupillary CV models (N = 1,408) control for age, sex, sampling rate, valid block count, and percent artifact; BCEA models (N = 2,238) control for the same covariates excluding percent artifact, which is undefined without a pupil signal. * q < 0.05 (FDR).

| Physiological Metric | Model Type | Predictor | $\beta$ | p | q (FDR) | $\Delta R^2$ |
| --- | --- | --- | --- | --- | --- | --- |
| Pupil Relative Volatility (CV) | Separate | SRS | -0.0187 | 0.4844 | 0.6458 | 0.0003 |
|  | Separate | SWAN | 0.0007 | 0.9786 | 0.9786 | 0.0000 |
|  | Joint | SRS | -0.0240 | 0.4234 | 0.5645 | 0.0004 |
|  | Joint | SWAN | 0.0121 | 0.6957 | 0.6957 | 0.0004 |
|  | Interaction | SRS x SWAN | -0.0646 | 0.0116 | 0.0233 * | 0.0045 |
| Spatial Gaze Instability (BCEA) | Separate | SRS | 0.0682 | 0.0004 | 0.0007 * | 0.0046 |
|  | Separate | SWAN | 0.0728 | 0.0002 | 0.0007 * | 0.0049 |
|  | Joint | SRS | 0.0445 | 0.0417 | 0.0833 | 0.0064 |
|  | Joint | SWAN | 0.0504 | 0.0259 | 0.0833 | 0.0064 |
|  | Interaction | SRS x SWAN | -0.0248 | 0.1771 | 0.1771 | 0.0007 |

For pupillary CV, neither SRS nor SWAN showed a significant main effect (all p ≥ .42; ΔR² ≤ 0.0004). A significant SRS × SWAN interaction emerged (β = −0.065, p = .012, q = .023, ΔR² = 0.0045). Joint elevation of autistic and ADHD trait severity was associated with disproportionately lower pupillary volatility than either trait alone, a sub-additive pattern paralleling the categorical interaction (Section 3.3). Given the small effect size, we interpret this as a modest but consistent signal rather than a robust dimensional marker.

For BCEA, separate models showed significant positive associations with SRS (β = 0.068, p = .0004, q = .0007) and SWAN (β = 0.073, p = .0002, q = .0007), attenuating but remaining nominally significant jointly (SRS: β = 0.045, p = .042; SWAN: β = 0.050, p = .026). A hardware-stratified check (Supplementary Section S2.1), however, showed these associations were absent at 30/60 Hz, the corneal-reflection-validated configurations, and significant only at 120 Hz, which lacks this validation and uses distinct onboard gaze processing. We therefore do not interpret the pooled BCEA-trait associations as genuine, attributing them instead to hardware-related acquisition differences that co-varied with sample composition across sites.

The SRS × SWAN interaction on BCEA was not significant (β = −0.025, p = .18).

Collectively, dimensional trait severity showed no hardware-independent association with gaze instability, and a modest sub-additive interaction effect on pupillary volatility (Supplementary Figure S1).

### 3.3. Categorical (DSM-5) Analysis of Clinical Diagnoses

A 2×2 factorial ANCOVA (ASD Status × ADHD Status) compared isolated diagnoses, comorbid presentations, and typically developing controls (baseline reference). Pupillary CV (N = 1,007) and BCEA (N = 1,633) models draw on different analytic samples and covariate sets (Table 3). Both metrics showed significant categorical main effects and a shared sub-additive comorbidity pattern, robust for CV and more equivocal for BCEA.

**Figure 1.**
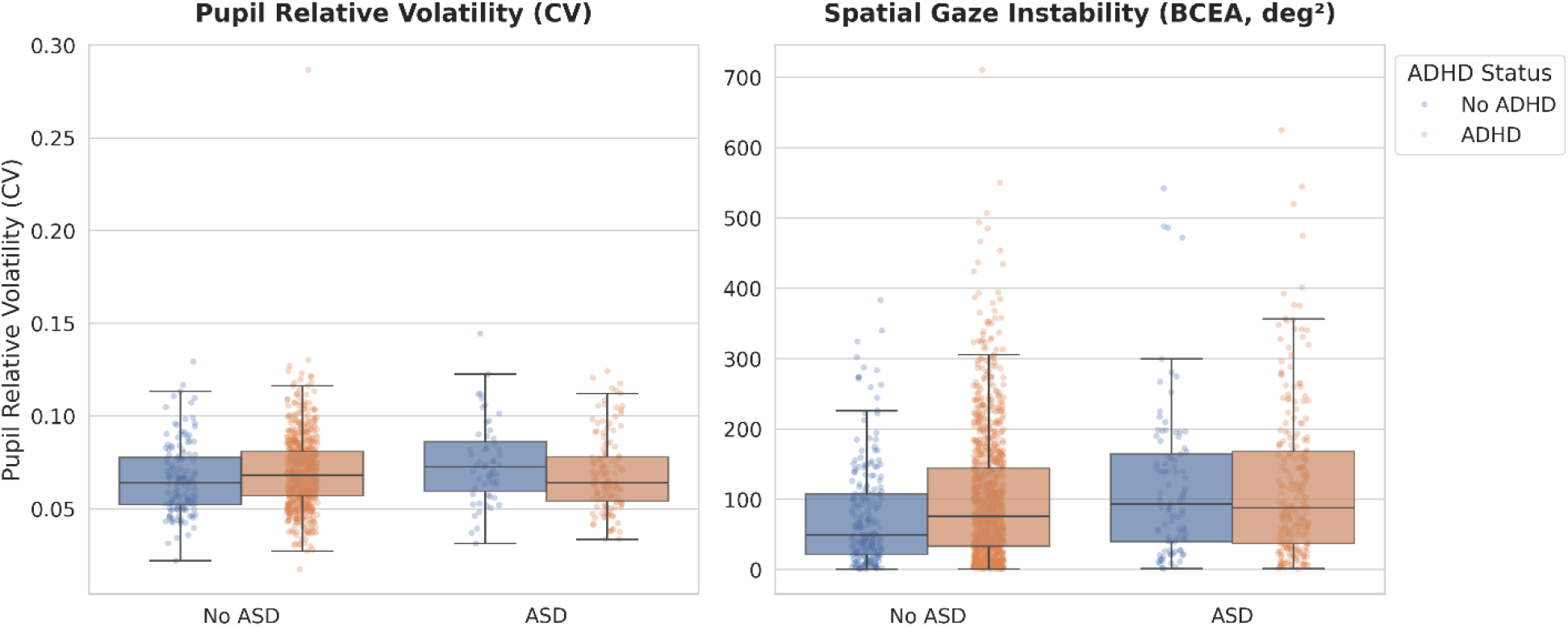
Categorical resting-state oculomotor physiology by DSM-5 diagnostic group. Box plots with individual data points (strip plots), 2×2 design comparing isolated diagnoses, comorbid presentations, and controls. (Left) Pupillary CV: significant ASD elevation and sub-additive comorbid interaction. (Right) BCEA: significant ASD and ADHD main effects, with a directionally consistent but estimation-sensitive comorbid interaction. Boxes show median/IQR; points are raw, unadjusted values. CV and BCEA panels reflect different analytic samples (N = 1,007, 1,633); adjusted models in Table 3.

**Table 3.** Categorical 2×2 ANCOVA models; typically developing controls are the baseline reference. Pupillary CV models (N = 1,007) control for age, sex, sampling rate, valid block count, and percent artifact; BCEA models (N = 1,633) control for the same covariates excluding percent artifact, which is undefined without a pupil signal. * q < 0.05 (FDR).

| Metric | Effect Term | $\beta$ | p | q (FDR) | $\eta_p^2$ |
| --- | --- | --- | --- | --- | --- |
| Pupil Relative Volatility (CV) | Main Effect: ASD | 0.4282 | 0.0077 | 0.0116 * | 0.0071 |
|  | Main Effect: ADHD | 0.1905 | 0.0421 | 0.0505 | 0.0041 |
|  | Interaction: ASD x ADHD | -0.5070 | 0.0059 | 0.0116 * | 0.0076 |
| Spatial Gaze Instability (BCEA) | Main Effect: ASD | 0.3902 | 0.0006 | 0.0034 * | 0.0073 |
|  | Main Effect: ADHD | 0.1821 | 0.0067 | 0.0116 * | 0.0045 |
|  | Interaction: ASD x ADHD | -0.2052 | 0.1105 | 0.1105 | 0.0016 |

#### 3.3.1. Pupillary Volatility

Isolated ASD diagnosis was associated with significantly elevated CV relative to controls (β = 0.4282, p = .0077, q = .0116); isolated ADHD showed no corresponding effect (β = 0.1905, p= .0421, q = .0505). A significant negative ASD × ADHD interaction (β = −0.5070, p = .0059, q = .0116) indicated that the elevation associated with ASD alone was largely offset when ADHD was also present. This parallels the modest sub-additive SRS × SWAN interaction observed dimensionally, suggesting comorbid autistic and ADHD-related features attenuate pupillary volatility whether indexed categorically or dimensionally.

#### 3.3.2. Gaze Instability

BCEA showed a convergent main-effect pattern: both isolated ASD (β = 0.3902, p = .0006, q = .0034) and isolated ADHD (β = 0.1821, p = .0067, q = .0116) were associated with significantly increased gaze instability relative to controls. The interaction was directionally consistent with CV’s (negative, sub-additive) but did not reach significance (β = −0.2052, p = .1105).

#### 3.3.3. Robustness of the Categorical Effects

All effects above were evaluated against outlier-resistant (robust regression), hardware, and medication sensitivity analyses (Supplementary Sections S1–S3; full statistics therein).

For pupillary CV, the ASD main effect and interaction were materially unchanged across all three checks. The pupil-CV-eligible sample is drawn almost entirely from the two hardware-consistent tracker configurations, so hardware heterogeneity does not threaten this result.

For BCEA, the ASD main effect was similarly robust throughout. The ADHD main effect held under robust regression and hardware stratification but attenuated below significance after medication adjustment (β = 0.1315, q = .0798); because stimulant use was concentrated in the ADHD and comorbid groups, this likely reflects the medication covariate partly absorbing ADHD-associated variance rather than indicating a pharmacological artifact, but we treat this effect with corresponding caution. The BCEA interaction remained non-significant under hardware stratification and medication adjustment, reaching only nominal significance under robust regression (β = −0.2240, p = .0311); given the modest shift from the non-robust estimate (−0.205 to −0.224), this most likely reflects reduced residual variance under M-estimation rather than correction of a biased coefficient, and we interpret this interaction cautiously throughout.

In sum, both metrics showed robust categorical main effects for isolated ASD (and, with the noted caveat, ADHD), with a sub-additive comorbidity interaction that was consistently robust for pupillary CV and directionally consistent but more estimation-sensitive for BCEA.

## 4. Discussion

The present study investigated intrinsic pupillary and oculomotor physiology across the ASD-ADHD spectrum during an unconstrained resting state in a large pediatric cohort, evaluated in parallel through dimensional and categorical frameworks. Contrary to our initial expectation that these frameworks would map onto distinct physiological metrics, categorical diagnostic status was the more consistent organizing factor for both pupillary CV and BCEA, each showing a directionally consistent sub-additive comorbidity interaction. Dimensional models showed no significant main effects for either metric, with one exception, a modest SRS × SWAN interaction on pupillary CV echoing the categorical comorbidity signature in continuous form. Reaching this picture required substantial methodological scrutiny: multi-site hardware heterogeneity produced an apparent dimensional BCEA association that did not survive validation, underscoring that transdiagnostic eye-tracking findings of this kind warrant explicit hardware-level verification before interpretation, a point we return to below.

### 4.1. Categorical Differentiation of Pupillary Volatility

Pupillary CV showed a robust categorical signature, an isolated-ASD main effect and a sub-additive ASD × ADHD interaction, both stable across every sensitivity analysis, despite the absence of any significant dimensional association with continuous trait severity. This divergence is worth taking at face value rather than explaining away: in this cohort, categorical diagnostic membership captured pupillary variance that continuous SRS-2/SWAN scores did not.

Spontaneous pupillary fluctuation during rest has been proposed as a peripheral correlate of autonomic tone, with mechanistic links to the LC-NE system established primarily for the phasic, stimulus-evoked response (Aston-Jones and Cohen, 2005; Joshi et al., 2016); the substrates of tonic, resting-state variability specifically remain less characterized, and we treat CV as a candidate correlate rather than a validated arousal biomarker. A recent task-evoked study found pre-saccadic pupil size relates to reaction time differently by diagnosis, linearly in autism, quadratically in ADHD (Bellato et al., 2023), consistent with diagnosis-specific rather than trait-graded arousal-oculomotor coupling, albeit under stimulus-driven conditions. Our findings extend this picture to an unconstrained baseline and to comorbidity specifically, while remaining agnostic about mechanism. This also diverges from prior work reporting continuous scaling of pupillary measures with autism trait severity (DiCriscio and Troiani, 2017); differences in paradigm, trait measure, and sample composition may account for this, but the discrepancy itself argues against treating either dimensional or categorical accounts of pupillary physiology as settled.

### 4.2. Categorical Differentiation of Gaze Instability

BCEA showed an even more uniformly robust categorical profile: significant main effects for both isolated ASD and isolated ADHD, stable across every sensitivity check. Increased gaze dispersion has recently been reported directly in ADHD using this same metric (Moiroud et al., 2025); our findings extend this to autism and to a large transdiagnostic sample at rest rather than during active fixation.

The BCEA interaction was directionally consistent with CV’s but reached significance only under robust regression, for reasons discussed above (Section 3.3.3); we treat it as suggestive and convergent with, but less certain than, the CV finding. Unlike CV, the pooled dimensional BCEA-trait association did not survive hardware-consistent restriction and is not interpreted as genuine (Section 3.2), a reminder that categorical and dimensional findings for the same metric can differ not only in strength but in validity, and each warrants independent scrutiny rather than a shared inference.

### 4.3. A Convergent Sub-Additive Comorbidity Signature

Beyond the individual main effects, both metrics converged on a common comorbidity pattern: the ASD+ADHD group did not show the cumulative elevation predicted by summing the isolated effects, but rather a negative interaction indicating attenuation relative to either condition alone. This was strongly supported for pupillary CV and directionally consistent but more estimation-sensitive for BCEA; we treat the two as converging evidence for a shared sub-additive effect of differing strength.

A non-additive ASD-ADHD relationship has been reported in other oculomotor and social-attentional measures, with gaze-cueing and intra-subject variability findings leading one group to conclude that comorbid presentations require distinct explanatory models rather than a combination of each condition’s isolated effects (Seernani et al., 2021). A conceptually related pattern emerged independently in a companion naturalistic movie-viewing investigation from our group using the same HBN cohort: comorbid ADHD paradoxically buffered the severe spatiotemporal gaze-phase decoupling otherwise associated with isolated ASD, localized specifically to social narrative contexts (Di and Biswal, 2026). Despite indexing different aspects of gaze physiology, dynamic synchrony during stimulus-driven viewing versus intrinsic fluctuation at rest, both converge qualitatively: comorbid ASD+ADHD produces an attenuated, non-additive phenotype rather than compounding either condition’s signature. This convergence across paradigms, measures, and analytic pipelines strengthens confidence that the sub-additive pattern is a genuine feature of comorbidity rather than a design-specific artifact.

The underlying mechanism cannot be determined from the present data. One plausible account is that isolated ASD and ADHD reflect different, partly opposing oculomotor and attentional regulatory profiles, for instance, heightened sensory reactivity versus impaired sustained attention, such that co-occurrence produces partial mutual offset rather than summation. We raise this as a hypothesis for future mechanistic work rather than a claim this design can adjudicate.

### 4.4. Methodological Considerations: Hardware Heterogeneity and Analytic Rigor

A central methodological finding of this study extends beyond either specific physiological metric: apparent dimensional associations in pooled, multi-site eye-tracking data can arise from acquisition heterogeneity rather than genuine trait-linked physiology, and this is not always captured by covarying for nominal sampling rate alone. One tracker configuration in this cohort lacked corneal-reflection-based pupil correction and provided no calibrated diameter output, while gaze calibration geometry varied at the session level rather than uniformly by hardware. Left unaddressed, this heterogeneity produced a significant pooled association between gaze instability and trait severity that did not survive restriction to hardware-consistent recordings and is not interpreted as genuine. By contrast, every categorical finding reported here remained stable across hardware-consistent subgroups, robust estimation, and medication adjustment — convergent evidence against an acquisition-driven artifact. Covariate adjustment for hardware is not equivalent to verifying that a signal is measured comparably across acquisition contexts in the first place. The two are easily conflated, particularly in large, open, multi-site pediatric datasets collected over extended periods whose published protocol documentation may not reflect every subsequent change in instrumentation. We encourage direct inspection of raw acquisition metadata in similar datasets rather than reliance on nominal covariates alone, particularly for dimensional or continuous findings.

The modest magnitude of the effects reported here (partial η² ≈ .005–.008; r ≈ .07–.09) merits similar interpretive care. Large-sample neuroimaging work indicates that small, reliably estimated effects, rather than the larger effects historically reported in small, underpowered studies, represent the genuine, reproducible signal (Marek et al., 2022), and effects of this magnitude are conventionally considered small at the level of a single observation but potentially consequential in aggregate rather than negligible (Funder and Ozer, 2019). Observed effect sizes are also mechanically constrained by measurement reliability, which has been identified as a major limiting factor in task-fMRI biomarker research specifically (Elliott et al., 2020; Vul et al., 2009). One recommended remedy is to increase within-subject sampling in order to stabilize individual-level estimates. Our own block-level, rather than pooled, computation of both metrics already reflects this principle, which may partly explain why this approach revealed associations that pooled estimation did not.

### 4.5. Limitations and Future Directions

The cross-sectional design precludes evaluating how these metrics change across development, and unconstrained resting-state eye-tracking remains sensitive to compliance and head movement despite our data-quality controls. The categorical analyses were affected by substantial group-size imbalance (isolated ASD n = 93 versus isolated ADHD n = 1,066), which may reduce precision for interaction-term estimation despite stability across sensitivity analyses. Complete daily medication records were unavailable for most participants, and missingness was associated with diagnostic group and autistic trait severity. Medication-adjusted models suggest this does not primarily drive our findings, but this could not be verified for the full sample. Categorical diagnoses were established via structured clinical interview rather than gold-standard instruments such as the ADOS/ADI, and the SRS-2 and SWAN, while validated trait measures, are not fully specific to ASD and ADHD in a community sample and may partly reflect broader psychopathology; this specificity concern applies most directly to the largely null dimensional findings and less to the categorical results, which rest on clinical consensus diagnosis rather than questionnaire threshold. Cognitive ability (FSIQ) was not included as a covariate, though HBN’s own IQ ≥ 70 inclusion criterion limits the range over which this could confound our findings; we consider this a reasonable target for confirmatory follow-up. Finally, as a community-referred rather than epidemiologically representative sample enriched for family-perceived clinical concern, generalizability to unselected populations should be considered with this ascertainment strategy in mind.

These findings also raise questions a resting-state design cannot answer alone, whether the sub-additive comorbidity signature is a fixed trait or context-dependent. Our companion naturalistic movie-viewing investigation in the same cohort begins to address this, finding that comorbid ADHD’s influence on ASD-associated gaze physiology is itself context-specific, emerging during social narrative viewing (Di and Biswal, 2026). Together, this suggests ASD-ADHD comorbidity may be best characterized not by a single physiological signature but by a consistent qualitative principle, non-additive, partially offsetting effects, that manifests differently by cognitive and sensory demand, a possibility future work directly contrasting resting and task-engaged physiology within the same individuals could test formally.

## 5. Conclusions

Categorical diagnostic status, rather than continuous trait severity, was the more consistent organizing factor for baseline pupillary and oculomotor physiology in this cohort. Both metrics showed a sub-additive comorbidity effect, most robustly for pupillary volatility, more provisionally for gaze instability, indicating that comorbid ASD and ADHD produce a distinct physiological profile rather than a simple combination of either condition’s isolated effects. Reaching this conclusion required explicit verification of acquisition hardware consistency alongside conventional statistical covariate adjustment, a distinction we suggest is broadly relevant to transdiagnostic research using large, open, multi-site pediatric datasets.

## Declarations

### Ethics approval and consent to participate

The data utilized in this study were obtained from the publicly available Healthy Brain Network (HBN) initiative. The original HBN study protocol was approved by the Chesapeake Institutional Review Board (now Advarra; Pro00012309), and all research was conducted in accordance with the Declaration of Helsinki. Prior to data collection, written informed consent was obtained from all adult participants and the legal guardians of all participants under 18 years of age. Written assent was additionally obtained from all participants under 18 years of age.

### Availability of data and materials

The datasets analyzed during the current study are publicly available through the Child Mind Institute’s Healthy Brain Network open-access initiative. The data can be accessed via the International Neuroimaging Data-sharing Initiative (INDI) at http://fcon_1000.projects.nitrc.org/indi/cmi_healthy_brain_network/.

## Abbreviations

ASD: Autism Spectrum Disorder
ADHD: Attention-Deficit/Hyperactivity Disorder
CV: Coefficient of Variation
BCEA: Bivariate Contour Ellipse Area
SRS: Social Responsiveness Scale
SWAN: Strengths and Weaknesses of ADHD symptoms and Normal behaviors
ANCOVA: Analysis of Covariance

