## Supplementary materials for "Sub-Additive Effects of Autism-ADHD Comorbidity on Resting-State Pupillary and Oculomotor Phenotypes"

This document provides supplementary analyses and data characterizations to support the primary findings reported in the main manuscript. The following sections evaluate the robustness of the observed resting-state physiological associations across alternative modeling approaches, hardware configurations, and potential clinical confounds.

- **Supplementary Figure S1** presents scatterplots of the dimensional associations between continuous autistic and ADHD trait severity and both physiological metrics, corresponding to the null findings reported in Section 3.2.
- **Section S1** addresses the influence of extreme values on the categorical diagnostic findings using robust regression (Huber's T M-estimation).
- **Section S2** evaluates the stability of the dimensional and categorical findings across the eye-tracking hardware configurations present in the dataset, given documented differences in pupil-channel calibration and gaze-signal processing between configurations (Section 2.4).
- **Section S3** details a sensitivity analysis regarding psychostimulant medication, including an assessment of data missingness, prevalence within the clinical subgroups, and a re-evaluation of the categorical models adjusting for medication status.

Together, these supplementary evaluations assess the stability of the reported resting-state physiological associations across alternative statistical approaches, hardware configurations, and clinical confounds.

**Supplementary Figure S1.** Dimensional associations between resting-state oculomotor physiology and transdiagnostic clinical traits. Scatter plots illustrating the continuous dimensional relationships between intrinsic eye-tracking metrics (x-axes) and behavioral symptom severity (y-axes) across the entire heterogeneous cohort. (Top row) Associations with continuous autistic traits (Social Responsiveness Scale; SRS-2 T-scores). (Bottom row) Associations with ADHD traits (SWAN Total scores). (Left column) Pupillary relative volatility (CV). (Right column) Spatial gaze instability (BCEA). Solid trend lines represent the unadjusted bivariate linear regression estimates, with shaded regions indicating 95% confidence intervals.

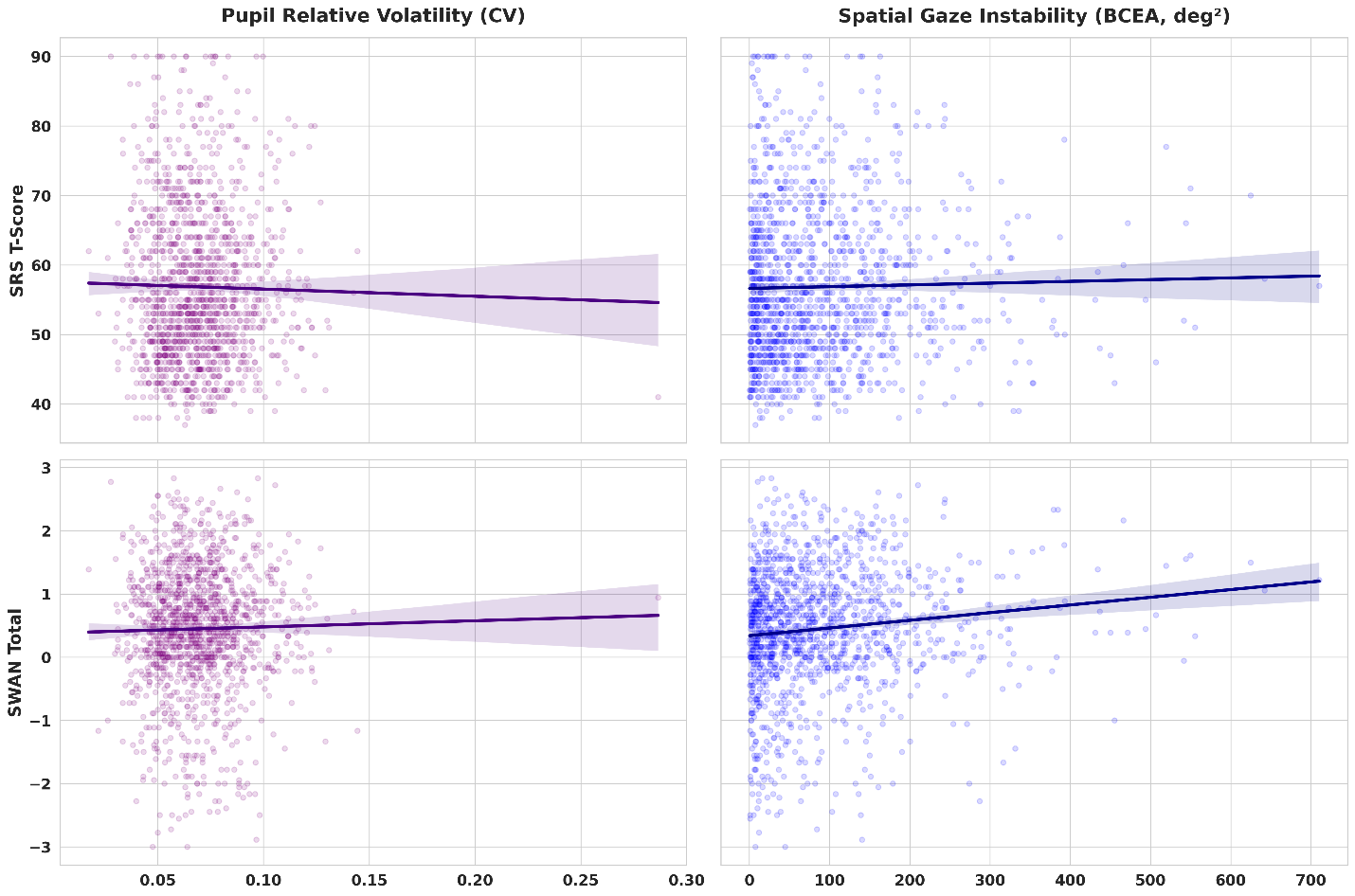

**S1. Robust Regression Sensitivity Analysis for Categorical Effects**

Both physiological metrics are variance-based and could in principle be sensitive to a small number of extreme values arising from hardware tracking artifacts, intrusive microsaccades, or excessive participant movement. To assess whether the categorical effects reported in Table 3 were disproportionately driven by such values, we re-evaluated the 2×2 categorical models for pupillary CV (N = 1,007) and BCEA (N = 1,633) using a Robust Linear Model (RLM) with Huber's T M-estimation, which mathematically down-weights the influence of extreme statistical residuals (Table S1).

For pupillary relative volatility (CV), all three effects observed in the primary model remained significant under robust estimation, with point estimates closely matching the original OLS results: the ASD main effect (β = 0.4162, p = .0053, q = .0080; cf. β = 0.4282 in Table 3), the ADHD main effect (β = 0.2068, p = .0176, q = .0211; cf. β = 0.1905, non-significant at q < .05 in Table 3), and the ASD × ADHD interaction (β = −0.5291, p = .0020, q = .0059; cf. β = −0.5070 in Table 3). The close correspondence in effect magnitude across estimation methods indicates these results are not driven by extreme residuals.

For spatial gaze instability (BCEA), the ASD and ADHD main effects were also confirmed under robust estimation (ASD: β = 0.3402, p = .0002, q = .0012; ADHD: β = 0.1604, p = .0031, q = .0062), with point estimates again closely matching the primary model. The ASD × ADHD interaction, which did not reach significance in the primary OLS model (β = −0.2052, p = .1105; Table 3), reached nominal significance under robust estimation (β = −0.2240, p = .0311, q = .0311). Because the point estimate itself shifted only modestly between methods (−0.205 to −0.224), this change in significance most likely reflects a reduction in estimated residual variance under M-estimation rather than correction of a substantively biased coefficient; consistent with the interpretation in Section 3.3.2, we treat the BCEA interaction as a directionally consistent but less certain effect rather than a confirmed finding on equal footing with the pupillary CV interaction.

**Supplementary Table S1.** Robust regression sensitivity analysis of categorical main effects and interactions. Standardized results from Robust Linear Models (RLM) using Huber's T M-estimation, testing the stability of the categorical DSM-5 diagnostic effects reported in Table 3 against extreme statistical residuals. Pupillary CV models (N = 1,007) adjust for participant age, biological sex, hardware sampling rate, number of valid eye-open data blocks, and the proportion of pre-interpolation flagged pupil samples per participant; BCEA models (N = 1,633) adjust for the same covariates with the exception of the pupil-derived flag-rate covariate, which does not apply to a gaze-based outcome (Section 2.4). * indicates significance at FDR-corrected q < 0.05.

| Physiological Metric | Effect Term | Beta | p-value | q-value (FDR) |
| --- | --- | --- | --- | --- |
| Pupil Relative  Volatility (CV) | Main Effect: ASD | 0.4162 | 0.0053 | 0.0080 * |
|  | Main Effect: ADHD | 0.2068 | 0.0176 | 0.0211 * |
|  | Interaction: ASD x ADHD | -0.5291 | 0.0020 | 0.0059 * |
| Spatial Gaze  Instability (BCEA) | Main Effect: ASD | 0.3402 | 0.0002 | 0.0012 * |
|  | Main Effect: ADHD | 0.1604 | 0.0031 | 0.0062 * |
|  | Interaction: ASD x ADHD | -0.2240 | 0.0311 | 0.0311 * |

**S2. Hardware Subgroup Sensitivity Analysis**

**S2.1. Dimensional Models**

To assess whether the dimensional associations reported in Section 3.2 were dependent on tracker hardware, we re-evaluated the separate SRS and SWAN models for pupillary CV and BCEA independently within each sampling-rate stratum (Table S2). For pupillary CV, neither predictor reached significance at 30 Hz (N=277) or 60 Hz (N=1,131; all q ≥ .58), consistent with the null pooled result (Section 3.2). For BCEA, the pooled association with SRS and SWAN was not present at 30 Hz (N=277) or 60 Hz (N=1,346; all q ≥ .43) and was significant only at 120 Hz (N=615; SRS: q=.118; SWAN: β=0.131, q=.033), the tracker configuration lacking corneal-reflection-based pupil correction and possessing distinct onboard gaze-signal processing (Sections 2.3–2.4). This pattern indicates the pooled BCEA dimensional association was not hardware-independent and is not interpreted as a genuine trait-linked effect (Section 3.2).

**Supplementary Table S2.** Dimensional analysis model within each sampling rate.

| Metric | Sample Rate | Predictor | β | p | q (FDR) | N |
| --- | --- | --- | --- | --- | --- | --- |
| Pupil Relative  Volatility (CV) | 30 Hz | SRS | 0.0231 | 0.7185 | 0.7983 | 277 |
|  | 30 Hz | SWAN | 0.0281 | 0.6658 | 0.7983 | 277 |
|  | 60 Hz | SRS | -0.0365 | 0.2822 | 0.5835 | 1131 |
|  | 60 Hz | SWAN | 0.0129 | 0.7120 | 0.7983 | 1131 |
| Spatial Gaze  Instability (BCEA) | 30 Hz | SRS | 0.0608 | 0.3317 | 0.5835 | 277 |
|  | 30 Hz | SWAN | -0.0593 | 0.3501 | 0.5835 | 277 |
|  | 60 Hz | SRS | 0.0072 | 0.8018 | 0.8018 | 1346 |
|  | 60 Hz | SWAN | 0.0447 | 0.1300 | 0.4333 | 1346 |
|  | 120 Hz | SRS | 0.0956 | 0.0237 | 0.1183 | 615 |
|  | 120 Hz | SWAN | 0.1307 | 0.0033 | 0.0332 | 615 |

**S2.2. Categorical Models**

The categorical 2×2 models reported in Section 3.3 were similarly re-evaluated within each hardware stratum (Table S3; pupillary CV's 120 Hz stratum was excluded, as this configuration lacks a valid calibrated pupil channel and contributes negligible pupil-CV-eligible data). For pupillary CV, the ASD main effect and ASD × ADHD interaction were directionally consistent and comparable in magnitude at 30 Hz (N=185; ASD β=0.68; interaction β=−0.87) and 60 Hz (N=822; ASD β=0.42; interaction β=−0.49), matching the pooled estimates (Table 3); neither stratum individually reached FDR significance, consistent with reduced power following stratification rather than an absent effect. For BCEA, the ASD and ADHD main effects were similarly consistent in direction and magnitude at 60 Hz (N=985) and 120 Hz (N=463), matching the pooled result; at 30 Hz (N=185), the ASD main effect remained directionally consistent, while the ADHD main effect was near zero (β=−0.08) and did not replicate, likely reflecting this stratum's limited size and sparse representation of typically developing controls (Section 2.3–2.4) rather than a genuine reversal. The BCEA interaction term did not reach significance in any individual hardware stratum, consistent with its status as the least certain of the reported categorical effects (Section 3.3.2).

**Supplementary Table S3.** Categorical 2×2 ANCOVA within each sampling rate.

| Metric | Sample Rate | Effect Term | β | p | q (FDR) | N |
| --- | --- | --- | --- | --- | --- | --- |
| Pupil Relative  Volatility (CV) | 30 Hz | Main Effect: ASD | 0.6782 | 0.1371 | 0.2056 | 185 |
|  | 30 Hz | Main Effect: ADHD | 0.4783 | 0.1565 | 0.2134 | 185 |
|  | 30 Hz | Interaction: ASD x ADHD | -0.8739 | 0.0761 | 0.1426 | 185 |
|  | 60 Hz | Main Effect: ASD | 0.4171 | 0.0171 | 0.0641 | 822 |
|  | 60 Hz | Main Effect: ADHD | 0.1787 | 0.0686 | 0.1426 | 822 |
|  | 60 Hz | Interaction: ASD x ADHD | -0.4908 | 0.0151 | 0.0641 | 822 |
| Spatial Gaze  Instability (BCEA) | 30 Hz | Main Effect: ASD | 0.3892 | 0.3903 | 0.4878 | 185 |
|  | 30 Hz | Main Effect: ADHD | -0.0832 | 0.8041 | 0.8616 | 185 |
|  | 30 Hz | Interaction: ASD x ADHD | -0.0208 | 0.9660 | 0.9660 | 185 |
|  | 60 Hz | Main Effect: ASD | 0.3699 | 0.0106 | 0.0641 | 985 |
|  | 60 Hz | Main Effect: ADHD | 0.2091 | 0.0076 | 0.0641 | 985 |
|  | 60 Hz | Interaction: ASD x ADHD | -0.2697 | 0.1067 | 0.1778 | 985 |
|  | 120 Hz | Main Effect: ASD | 0.4570 | 0.0408 | 0.1225 | 463 |
|  | 120 Hz | Main Effect: ADHD | 0.2792 | 0.0698 | 0.1426 | 463 |
|  | 120 Hz | Interaction: ASD x ADHD | -0.1990 | 0.4247 | 0.4900 | 463 |

**S3. Psychostimulant Medication and Missing Data Characterization**

Due to the potential influence of ADHD psychostimulant medication on arousal and oculomotor dynamics, we characterized daily medication logs available within the Healthy Brain Network dataset. Of the 2,238 participants included in the BCEA dimensional models (Section 3.2; requiring complete SRS/SWAN data), daily medication logs were available for 467 participants.

Missing data were not Missing Completely at Random (MCAR). There were no significant differences in age (t = −0.95, p = .3417) or biological sex (χ² = 2.23, p = .1349) between those with and without medication data. However, completion of medication logs was significantly associated with clinical diagnostic grouping (χ² = 39.44, p < .0001), and parents of children with higher autistic traits were significantly more likely to complete these records (SRS-2 T-scores: M = 58.75 vs. 56.97, t = 2.77, p = .0057), whereas continuous ADHD traits did not significantly influence missingness (SWAN Total: t = −1.11, p = .2657) (Supplementary Table S4). The medication data are therefore considered Missing At Random (MAR), as missingness is systematically related to observed clinical variables (diagnosis and SRS-2 scores) already accounted for in our primary statistical models.

**Supplementary Table S4.** Comparison of participants with and without daily medication logs.

| Feature | Med Data  Available (n=467) | Med Data  Missing (n=1,771) | Test Statistic | p |
| --- | --- | --- | --- | --- |
| Age (Mean) | 10.83 | 10.99 | t=−0.95 | 0.3417 |
| Sex (% Female) | 40.3% | 36.4% | χ2=2.23 | 0.1349 |
| Diagnosis Group | See Table S5 | See Main Text | χ2=39.44 | < 0.0001 |
| SRS Total T-score | 58.75 | 56.97 | t=2.77 | 0.0057 |
| SWAN Total Score | 0.41 | 0.48 | t=−1.11 | 0.2657 |

Among the 467 participants with verified records, active stimulant use at the time of testing was recorded in 39 participants, representing an overall prevalence of 1.74% across the full analytic cohort (Supplementary Table S5). Within the isolated ADHD group, active stimulant use was 14.12%, and within the comorbid ASD+ADHD group, 17.31%. No active stimulant use was recorded in the typically developing control group or the isolated ASD group.

**Table S5.** Active Stimulant Use by Diagnostic Group.

| Diagnosis | No Stimulant | Taking Stimulant | % Taking Stimulant |
| --- | --- | --- | --- |
| ADHD without ASD | 152 | 25 | 14.12% |
| ASD and ADHD | 43 | 9 | 17.31% |
| ASD without ADHD | 21 | 0 | 0.00% |
| TD | 76 | 0 | 0.00% |
| Other | 136 | 5 | 3.55% |
| Total | 428 | 39 | 8.35% |

To evaluate whether medication status confounded our physiological findings, we re-ran the categorical 2×2 models, explicitly adding medication status (0 = unmedicated, 1 = active stimulant use, 2 = missing medication data) as a categorical covariate alongside the covariate sets described in Table 3 (Section 2.6.2), separately for the pupillary CV (N = 1,007) and BCEA (N = 1,633) analytic samples (Supplementary Table S6).

**Supplementary Table S6.** Categorical 2×2 ANCOVA sensitivity models adjusting for psychostimulant medication status. * indicates significance at FDR-corrected q < 0.05.

| Metric | Effect Term | Beta | p | q (FDR) | η_p_^2^ |
| --- | --- | --- | --- | --- | --- |
| Pupil Relative  Volatility (CV) | Main Effect: ASD | 0.4152 | 0.0096 | 0.0206 * | 0.0067 |
|  | Main Effect: ADHD | 0.1571 | 0.0965 | 0.1158 | 0.0028 |
|  | Interaction: ASD x ADHD | -0.4720 | 0.0103 | 0.0206 * | 0.0066 |
| Spatial Gaze  Instability (BCEA) | Main Effect: ASD | 0.3675 | 0.0011 | 0.0067 * | 0.0065 |
|  | Main Effect: ADHD | 0.1315 | 0.0532 | 0.0798 | 0.0023 |
|  | Interaction: ASD x ADHD | -0.1755 | 0.1709 | 0.1709 | 0.0012 |

The inclusion of the medication covariate largely confirmed the primary categorical findings. For pupillary CV, both the ASD main effect (β=0.4152, q=.0206) and the ASD × ADHD interaction (β=−0.4720, q=.0206) remained significant after pharmacological adjustment, closely matching the primary model (Table 3: β=0.4282, β=−0.5070). For BCEA, the ASD main effect remained significant (β=0.3675, q=.0067), also closely matching the primary model (β=0.3902), and the interaction remained non-significant, consistent with its status elsewhere in this paper (Sections 3.3.2, S1). The one effect not fully replicated was the BCEA ADHD main effect, which was significant in the primary model (Table 3: β=0.1821, q=.0116) but did not reach significance after medication adjustment (β=0.1315, q=.0798); given that active stimulant use was concentrated in the ADHD and comorbid groups (Table S5), this attenuation may reflect the medication covariate absorbing some of the variance associated with ADHD diagnosis itself, rather than indicating the original effect was a pharmacological artifact per se. We note this as a point of caution for the BCEA ADHD main effect rather than treating it as fully confirmed.
